# Transfer RNA modifications during rhizome development in *Oryza longistaminata*

**DOI:** 10.1101/2025.09.10.675464

**Authors:** Wenze Li, Guangzhao Yang, Haoyue Xue, Ruichen Ma, Chaoying Zhang, Ruixuan Yao, Yajun Li, Xukai Li, Fengyi Hu, Peng Chen, Zheng Li

## Abstract

The rhizome organ endows many clonal plants with the ability to reproduce under stressful climates. Such rhizome-mediated perenniality also underlies the recent success of perennial rice breeding. Despite the importance of this organ, its developmental regulation is poorly understood. Here, using *Oryza longistaminata*, an emerging model for rhizome biology, we explored the involvement of tRNA modification in rhizome development. Through comparative profiling of modified nucleotides in rhizome samples at different developmental stages and/or subjected to various treatments, we identified a number of key rhizome-related tRNA modifications and revealed that, akin to the observations in many non-plant organisms, tRNA modifications are associated with hormonal signalling to regulate rhizomes’ responses to environmental cues. Furthermore, genome-wide analysis of tRNA-modifying genes was performed to facilitate future functional studies. These results and analyses not only deepen our understanding of rhizome development but also underscore the regulatory roles of tRNA modification in plant organ development.

## BACKGROUND

Transfer RNA (tRNA), an essential class of small non-coding RNAs typically containing 70– 100 nucleotides, is responsible for carrying specific amino acids to the growing polypeptide chain by matching its anticodon to the mRNA codon. To ensure accurate and efficient protein synthesis, tRNAs are enzymatically modified post-transcriptionally (El Yacoubi et al. 2012). Compared with other RNA classes (e.g., mRNA and rRNA), modifications on tRNAs are much more extensive—around 10–20% of their nucleotides are modified, and a vast majority of RNA modification types we know so far have been identified in tRNAs (Roundtree et al. 2017). Research on humans revealed that a human tRNA molecule contains 13 modifications on average, with a great deal of variation in modification type and modification site (Saikia et al. 2010). Such a rich diversity of tRNA modification implicates its importance beyond protein synthesis. Indeed, recent studies in yeast and animals elucidated that deposition of modifications into tRNAs is a highly dynamic process, which responds to both environmental and developmental cues, and that these modifications may be involved in the regulatory networks of multiple cellular processes via modulating gene expression (Schimmel 2018; Frye et al. 2018). Notably, outstanding connections have now been established between deficiencies in tRNA modification and many human diseases, such as mitochondrial diseases, neurological disorders and cancer, sparking widespread research interest among the biomedical research community (Barbieri and Kouzarides 2020; Suzuki 2021).

In sharp contrast to such a surge of attention, research on tRNA modification in plants is still in its infancy. In our previous work, we for the first time established a high-performance liquid chromatography (HPLC)-based pipeline for profiling modified tRNA nucleosides in plants and discovered that there is also a wide range of modifications present in tRNAs from the model plant species *Arabidopsis* and hybrid aspen (Chen et al. 2010). Subsequently, using an improved liquid chromatography combined with tandem mass spectrometry (LC-MS/MS)-based approach, we systematically investigated the composition and abundance of tRNA modifications in rice and *Arabidopsis*, and revealed that they vary dramatically in response to various environmental stimuli and during different physiological conditions (Wang et al. 2017c). In a more specific study, Janssen et al. (2022) focused on the wobble position of *Arabidopsis* tRNAs and demonstrated that salt stress conditions can promote the occurrence of some specific modifications on this site. Consistent with these findings, mutants of genes encoding tRNA modifying enzymes were found to display defects in multiple developmental processes and show compromised stress tolerance (Nakai et al. 2019; Guo et al. 2019), and conversely, overexpression of certain such genes can lead to enhanced stress resistance mainly via regulating hormonal signalling pathways (Wang et al. 2017a). These molecular insights not only substantiate the regulatory significance of tRNA modification in plant physiology but also bear practical value for improving agricultural production. However, research on tRNA modification has largely been only conducted in a few model plant species, and its involvement in tissue-or organ-specific development has also not been explored.

Rhizomes are modified stems growing horizontally under the ground for vegetative reproduction. While this organ is absent in many major crop plants, it is of both agricultural and ecological importance (Guo et al. 2021). First, as rhizomes are usually rich in nutrients and secondary metabolites, they are the edible part of some rhizome-forming crops (e.g., lotus) and widely used in traditional Chinese medicine (Koo et al. 2013; Yang et al. 2015). On the other hand, rhizome-mediated vegetative reproduction confers aggressiveness to many invasive species, presenting a threat to biodiversity and agricultural production (Song et al. 2013). Despite these significant points, rhizomes attracted little attention until very recently, when rhizomatousness-associated perenniality has been exploited to perennialise annual rice cultivars (Zhang et al. 2023). Due to the multidimensional benefits of sustainability associated with perennial grain crops, their successful breeding is considered a milestone achievement in agricultural research. On this account, the necessity of understanding rhizome development and its regulatory mechanisms becomes evident.

As the rhizomatous parent of perennial rice breeding, *Oryza longistaminata* has emerged as a model for studying rhizome development (Li et al. 2022; Zhang et al. 2023). Previous research on this African wild rice species revealed that rhizomes generally experience two stages during their development. First, after a nascent rhizome forms from the stem base of the mother plant, its apical meristem keeps generating new nodes for elongation growth (Lian et al., 2024). When a rhizome is sufficiently elongated, its tip can bend upwards and subsequently develop into a new ramet above the ground, a process known as developmental transition (Yoshida et al., 2016). Interestingly, the occurrence of such a transition was found to be correlated with the total number of internodes rather than the absolute length of the rhizome, indicating that the elongation procedure of a rhizome can be roughly divided into different sub-stages according to the number of internodes (Bessho-Uehara et al. 2018). To date, only a few environmental and hormonal factors have been associated with rhizome development. For the elongation stage, nitrogen nutrition status has been found to control rhizome bud activity via regulating the cytokinin pathway (Shibasaki et al. 2021). Regarding the developmental transition stage, light exposure and the phytohormone gibberellin (GA) play a key role in influencing the shooting of rhizomes, and sucrose appears to be a crucial signalling compound, whose depletion initiates the transition (Koo et al. 2013; Bessho-Uehara et al. 2018).

In this study, to examine how tRNA modifications are involved in rhizome development in *O. longistaminata*, we systematically profiled tRNA modifications during both the elongation growth and developmental transition stages. By treating rhizomes with various environmental and hormonal factors that are known to affect rhizome development, we identified a number of key rhizome development-related tRNA modifications. These findings, together with genome-wide analysis of *O. longistaminata* genes encoding putative tRNA modification enzymes, highlight tRNA modification as an important layer of plant developmental regulation and offer added molecular insights into rhizome development.

## RESULTS

### Global tRNA modification profile of the O. longistaminata plant

To ensure that our tRNA modification profiling pipeline is suitable for *O. longistaminata* and also to obtain an overview of tRNA modifications in this plant, we first applied our LC-MS/MS-based approach to quantitatively analyse tRNA modifications in four *O. longistaminata* tissues, namely, the leaf, stem base, nascent tiller, and rhizome (Figure 1). In total, we were able to detect 32 types of tRNA modification events. Of these modifications, the majority were modifications in the four principal RNA bases, with 10 for uracil (U), 9 for adenine (A), 5 for guanine (G), and 4 for cytosine (C). Moreover, there were 23 methylation-related modifications (designated by ‘m’ in the symbol), accounting for approximately 72% of the total. Across different tissues, the identified modifications were conservative in terms of their types, with only the mnm^5^U modification being below the detection level in the rhizome. Regarding the abundance of tRNA modification, different tissues showed a great deal of variation. For example, τm^5^s^2^U, m^2^A, and ms^2^io^6^A were of a much higher abundance in the leaf than in the other tissues, while the m^2^_2_G modification was more pronouncedly deposited in the rhizome and stem base. These results indicate that tRNA modification events are highly tissue-specific in *O. longistaminata* and imply that modification profiles in different tissues may have varying responses to environmental stimuli, thereby prompting us to further investigate tRNA modifications in the rhizome, an organ that is not present in well-studied model plants.

**Figure 1.**
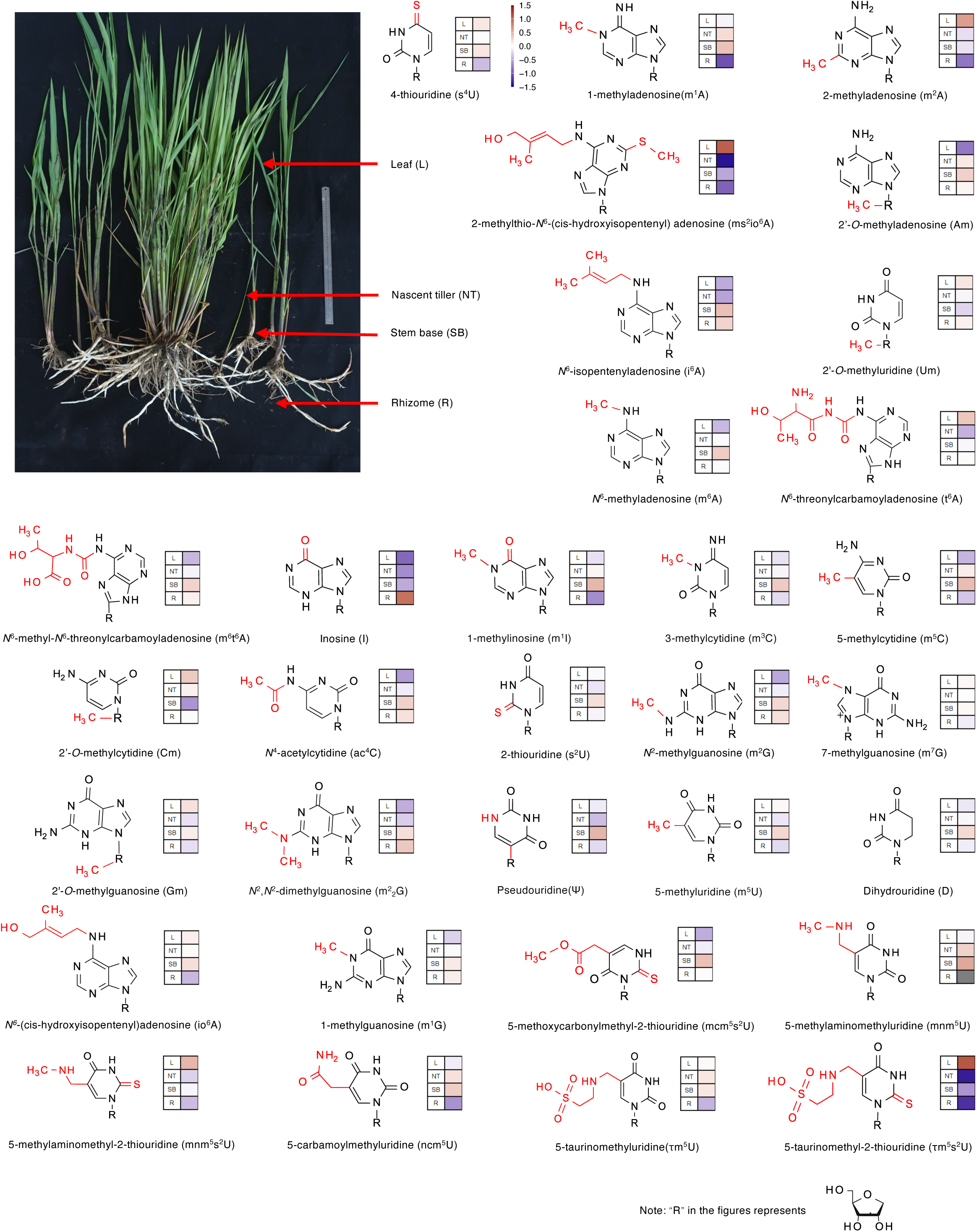
Global tRNA modification profile of the *O. longistaminata* plant. Leaf, nascent tiller, stem base and rhizome tissues were collected to perform tRNA profiling using liquid chromatography combined with tandem mass spectrometry (LC-MS/MS). The abundance of each modification was calculated relative to all detected nucleosides and normalised using into Z-score. The heatmap shows Z-score normalised data of tRNA abundance.

### Key tRNA modifications involved in the elongation stage of rhizome development

For the elongation stage of rhizome development, we took both the developmental and environmental cues into consideration, and two-factor ANOVA was used to identify important tRNA modifications during this stage. Specifically, rhizome samples were divided into three sub-stages according to the number of nodes (early stage: 1–2 nodes, middle stage: 3–5 nodes, and late stage: 6–8 nodes) (Figure 2A), and nitrogen nutrition, a soil condition that is known to influence rhizome development, was factored in by setting up two nitrogen treatments (High-N and Low-N).

**Figure 2.**
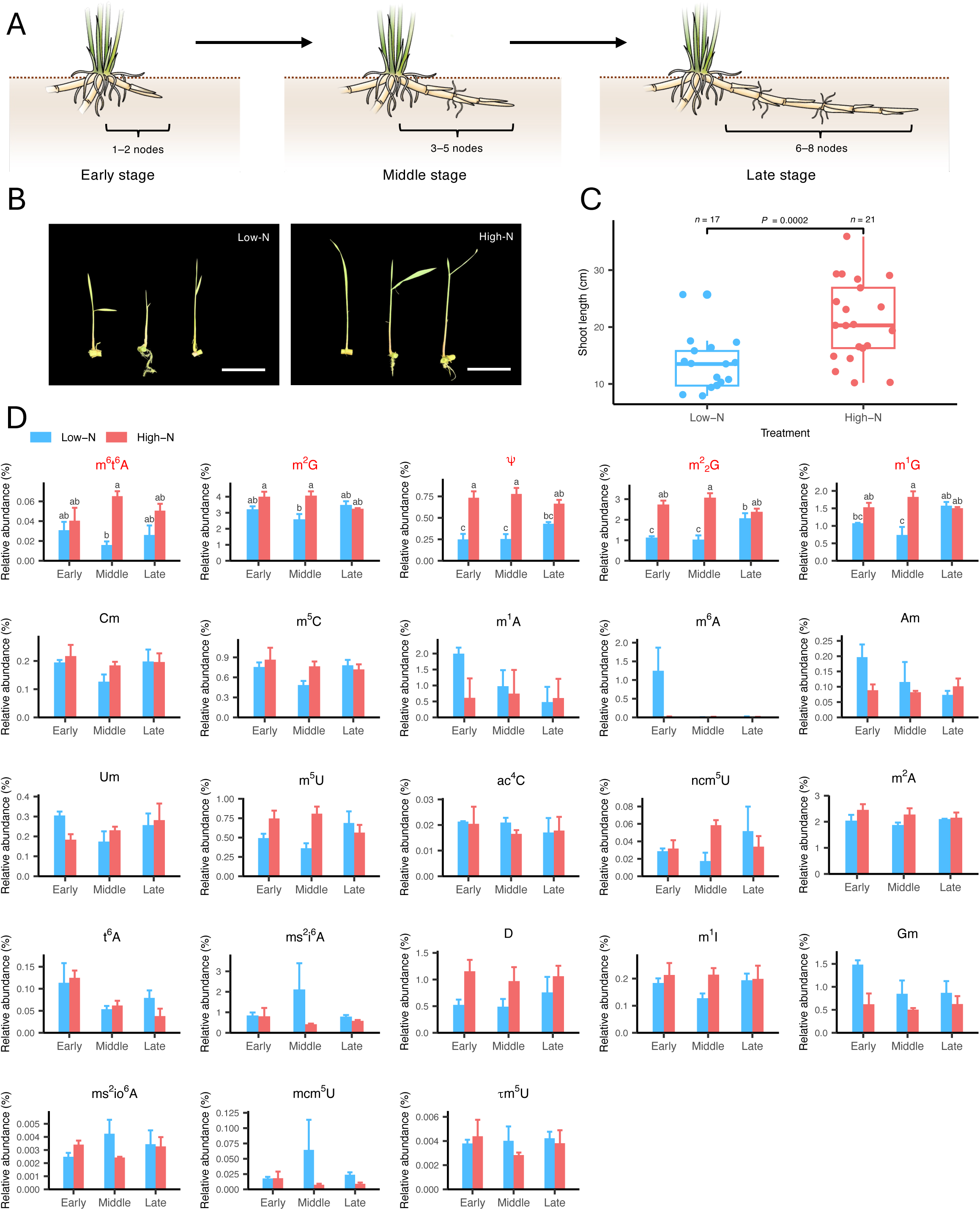
Key tRNA modifications involved in the elongation stage of rhizome development. **A** Schematic representations of rhizome development during the elongation stage, showing the three sub-stages of this elongation growth defined in this study. **B** Representative phenotype images showing the bud activity of rhizomes in the high-N and low-N groups. Bars = 5 cm. **C** Quantification of rhizome bud activity of the high-N and low-N groups. *P*-value was calculated using Student’s *t*-test. **D** tRNA modification profiles of rhizomes at three elongation sub-stages and subjected to high-N and low-N treatments. The abundance of each modification was calculated relative to all detected nucleosides. Data are displayed as mean ± SEM, n = 3. Significant differences were calculated by two-way analysis of variance followed by Tukey’s honest significant difference test. Compact letter displays were only shown for modifications that were detected to exhibit significant differences (*P* < 0.05; with the modification name highlighted in red).

We first confirmed the effect of nitrogen nutrition on rhizome growth and development by comparing bud activity of high-N and low-N treated rhizomes. Consistent with previous findings (Shibasaki et al. 2021), the high-N group displayed significantly higher bud activity in comparison with the low-N treatment (Figures 2B and 2C). Using modified Yoshida solutions with these two nitrogen concentrations, rhizomes at various developmental sub-stages were hydroponically cultured for two weeks and then subjected to tRNA modification profiling analysis (Figure 2D). Of all the detected modifications, m^2^G, m^2^_2_G, m^1^G, m^1^A, and m^2^A were generally of high quantity in all sample groups, with an average relative abundance ranging from 0.5% to 4.1%. These highly abundant modifications, however, did not all show significant level changes in our ANOVA results: two adenine-derived nucleosides (m^1^A and m^2^A) were not detected to be significantly differentially accumulated across developmental stages or under distinct nitrogen treatments. The other three rhizome abundant modifications (m^2^G, m^2^ G, and m^1^G), together with m^6^t^6^A and ψ, were identified as differentially behaving modifications. Notably, all these five modifications showed significantly increased enrichment upon elevated nitrogen application when rhizomes were at the middle developmental stage, indicating that rhizomes at this stage may be more responsive to environmental alterations. In addition to nitrogen nutrition, the factor of developmental stage also exerted a profound impact on m^2^_2_G and m^1^G, whose abundances were significantly promoted when low-N treated rhizomes elongated from the middle stage to the late stage.

### Key tRNA modifications involved in the developmental transition stage of rhizome development

One characteristic phenotypic feature of rhizomes undergoing developmental transition is upward bending at the far end, which is followed sequentially by emergence from the ground and development into an aerial stem (Figure 3A). Hence, to look into the developmental transition stage, we used bending angle change as a proxy for transition progression. As shown in Figures 3B and 3C, rhizomes that were dark cultured in 1/2 MS liquid media formed curvature with an angle of approximately 35 degrees after eight days, whereas such negative gravitropic bending was greatly repressed by sucrose supplement, and, to a lesser degree, by light exposure. These results indicate that both sucrose and light signals function as negative regulators of rhizomes’ developmental transition.

**Figure 3.**
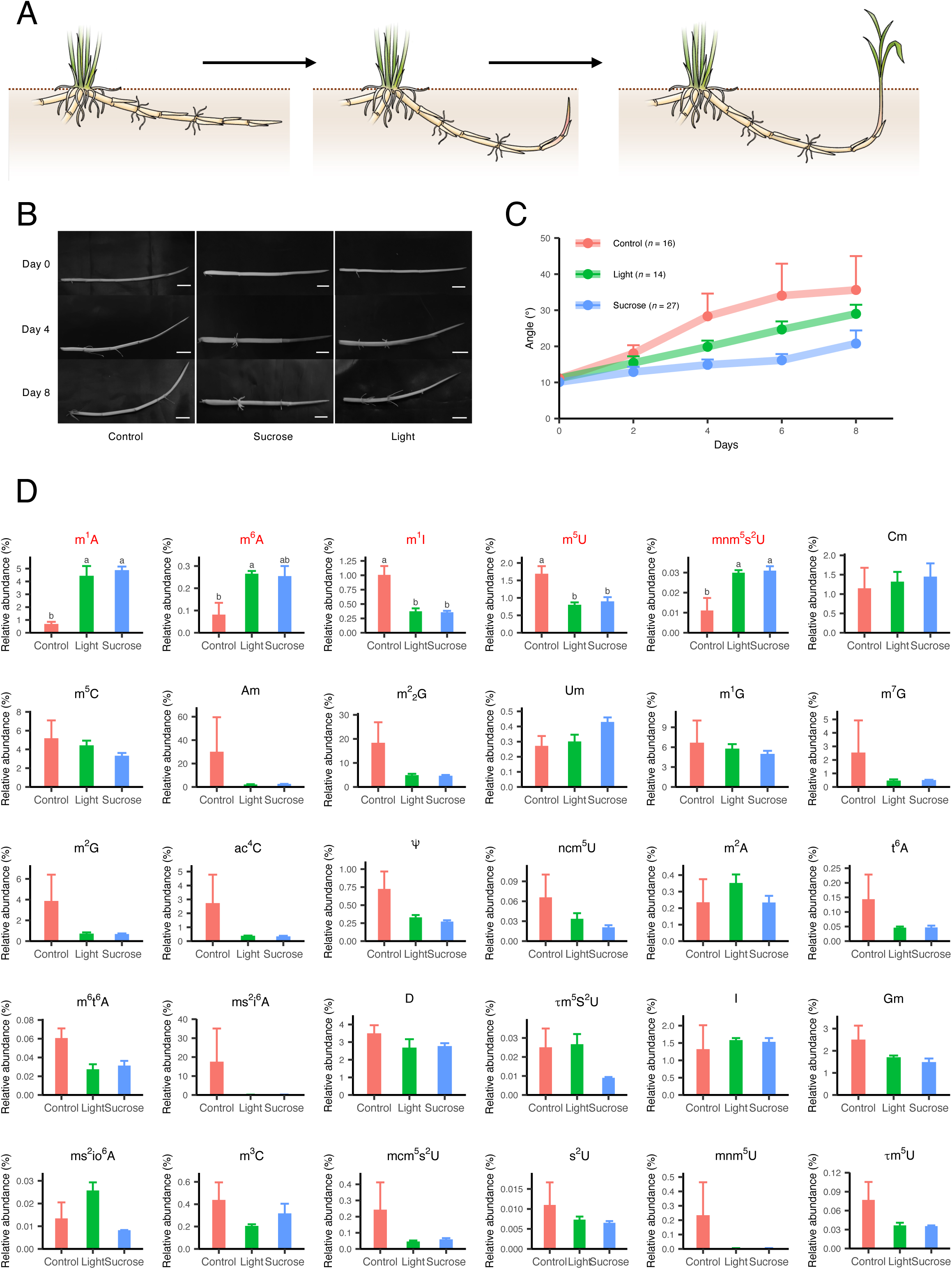
Key tRNA modifications involved in the developmental transition stage of rhizome development. **A** Schematic representations of rhizome development during the developmental transition stage. **B** Representative phenotype images showing the curvature growth of rhizomes in the light-treated, sucrose-treated, and control groups. Bars = 2 cm. **C** Quantification of bending angle changes of rhizomes in the light-treated, sucrose-treated, and control groups. Data are displayed as mean ± SEM. **D** tRNA modification profiles of rhizomes in the light-treated, sucrose-treated, and control groups. The abundance of each modification was calculated relative to all detected nucleosides. Data are displayed as mean ± SEM, n = 3. Significant differences were calculated by two-way analysis of variance followed by Tukey’s honest significant difference test. Compact letter displays were only shown for modifications that were detected to exhibit significant differences (*P* < 0.05; with the modification name highlighted in red).

Following these phenotypical observations, we excised the bending region of the rhizome from the aforementioned treatments and conducted tRNA nucleoside analysis (Figure 3D). Except for m^2^A, all the highly abundant modifications during the elongation stage (m^1^A, m^2^G, m^2^_2_G, and m^1^G) generally increased in abundance at the developmental transition stage, and some additional modified nucleosides were also enriched during this stage, such as m^5^C, D, and Gm, suggesting more intense involvement of tRNA modification during this rhizome-to-stem identity shift. Surprisingly, none of the key modifications identified during the elongation stage displayed significant quantitative variances under different culturing conditions, although methylation seemed to be important at both stages. Differential abundances were found between the treatments and the control group in another set of methylation-containing modifications (m^1^A, m^6^A, m^1^I, m^5^U, and mnm^5^s^2^U), and their patterns generally agreed with the phenotypes—the levels of m^1^I and m^5^U were significantly suppressed by the light and sucrose treatments, whereas these two conditions largely elevated m^1^A, m^6^A, and mnm^5^s^2^U, suggesting that these modification events are likely involved in the developmental transition of rhizomes with either a positive or a negative regulatory role.

### tRNA modifications in hormonal responses associated with rhizome development

Previous research has shown that tRNA modification may act in concert with hormonal signalling to control developmental responses to alterations of environmental conditions (Wang et al. 2017b; Chen et al. 2019). To understand how this action of mode functions in rhizome development, we treated plants with exogenous hormones that have known regulatory roles for the elongation and developmental transition stages, and compared their tRNA modification profiles with environmental factor-regulated profiles.

For the elongation stage, rhizome bud activity is influenced by the nitrogen level through regulation of cytokinin biosynthesis (Shibasaki et al., 2021). Indeed, rhizomes treated with 1 μM *trans*-zeatin tZ for two weeks exhibited promoted bud activity, a phenotype that mirrored those of rhizomes subjected to high-N treatment (Figure 4A). Despite these similar effects on the phenotype, tZ treatment led to significant abundance change in a markedly larger number of tRNA modifications, with a total of 20 modifications being either significantly induced or repressed (Figure 4B). Akin to the promotive effects of high-N conditions on certain tRNA modifications, a prominent increase of modified nucleosides was observed for all the tZ-induced differentially behaving modifications except for m^2^A and m^2^G. It is, however, noteworthy that such a level increase was detected often across the three developmental stages, rather than predominantly at the middle stage as for high-N induced differential modifications. Furthermore, the extent of exogenous cytokinin treatment’s effects was also more pronounced than those of high-N conditions, as demonstrated by multiple times of level increase in modifications such as τm^5^U, mnm^5^s^2^U, m^3^C, I, D, and m^7^G. Given the similar phenotypes resulting from the two treatments, we hypothesised that their shared differentially behaving modifications would be tRNA modifications relevant to the hormonal signalling pathway underpinning rhizome development. As shown in Figure 4C, we found three methylguanosine nucleosides (m^2^G, m^1^G, and m^2^ G) belonging to this category of critical modifications.

**Figure 4.**
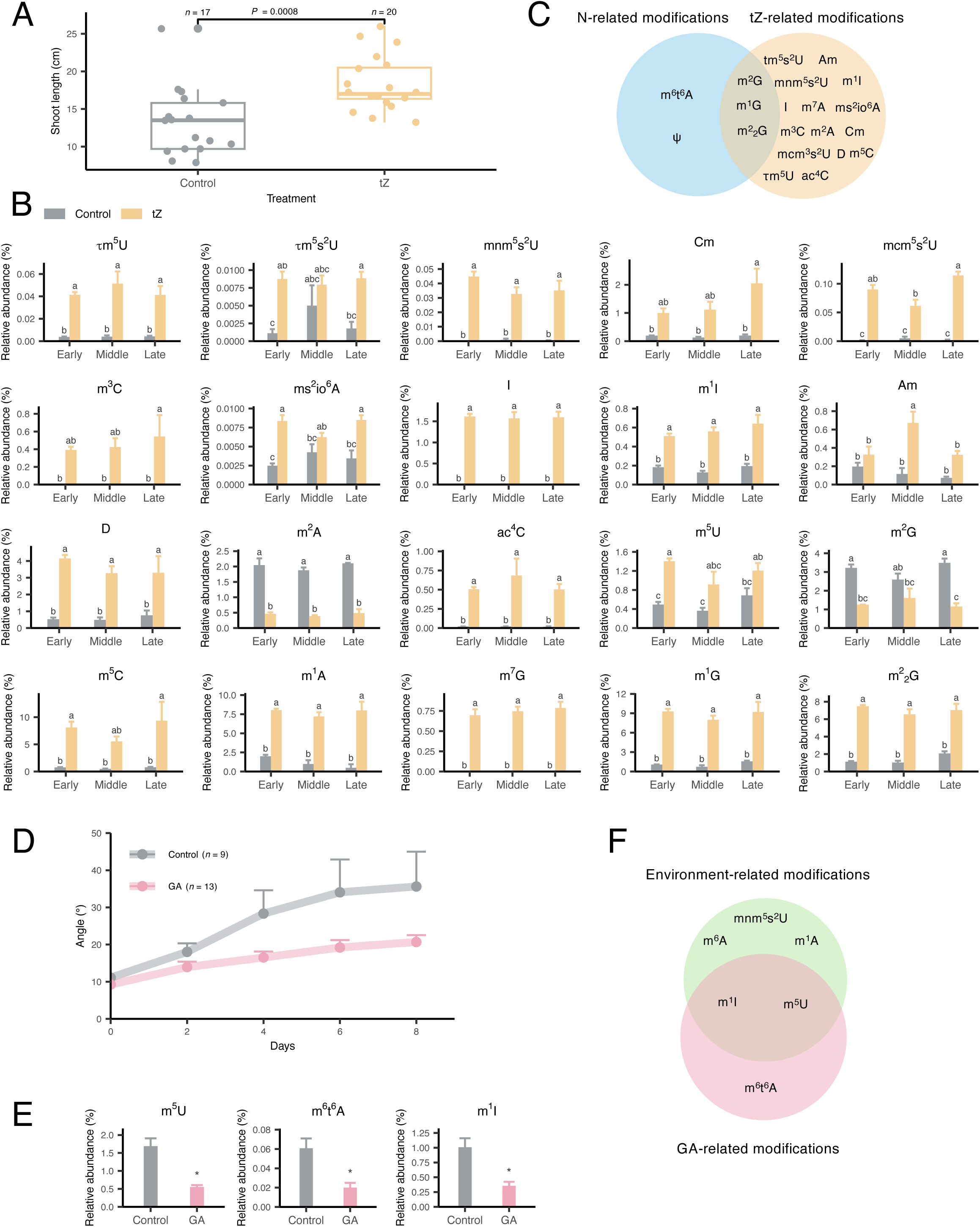
tRNA modifications in hormonal responses associated with rhizome development. **A** Rhizome bud activity of the control and tZ-treated groups. *P*-value was calculated using Student’s *t*-test. **B** tRNA modification profiles of rhizomes at three elongation sub-stages and/or subjected to tZ treatment. **C** Venn diagram showing unique and shared differentially behaving modifications detected in the nitrogen treatment and tZ treatment experiments. **D** Bending angle changes of rhizomes in the gibberellin (GA)-treated and control groups. Data are displayed as mean ± SEM. **E** tRNA modification profiles of rhizomes in the GA-treated and control groups. **F** Venn diagram showing unique and shared differentially behaving modifications detected in the environmental factor (sucrose and light) treatment and tZ treatment experiments. In **B** and **E**, the abundance of each modification was calculated relative to all detected nucleosides. Data are displayed as mean ± SEM, n = 3. Significant differences were calculated by two-way analysis of variance followed by Tukey’s honest significant difference test. Only modifications exhibiting significant differences are shown (*P* < 0.05).

In many plant species, the connection between GA and gravitropism is well established (Claeys et al. 2014). Thus, we assessed GA-induced tRNA modification abundance changes during the rhizome developmental transition stage. Compared with the control group, rhizomes *ex vivo* cultured in GA-containing medium exhibited suppressed negatively gravitropic responses, parallelling the phenomena observed for light-or sucrose-treated rhizomes (Figure 4D). Unlike the large-scale tRNA modification abundance changes stemming from exogenous cytokinin application, GA only significantly affected three modifications—m^6^t^6^A, m^5^U, and m^1^I (Figure 4E). Notably, the latter two of these modifications were shared differentially behaving modifications with the above experiment concerning environmental factors, and in both cases, modification levels were significantly inhibited in the treatment group (Figures 4E and 4F).

### Genome-wide analysis of tRNA modification-related genes in O. longistaminata

To probe molecular insights of tRNA modification during rhizome development, we next performed genome-wide analysis of tRNA modification-related genes in the *O. longistaminata* genome. By using protein sequences of genes responsible for tRNA modification in *S. cerevisiae* and *E. coli* (Chen et al. 2010), we identified a total of 76 homologous genes in *O. longistaminata* (Table 1). Except for m^6^A, *O. longistaminata* candidate genes were obtained for all differentially behaving modifications identified during the two developmental stages.

**Table 1.**
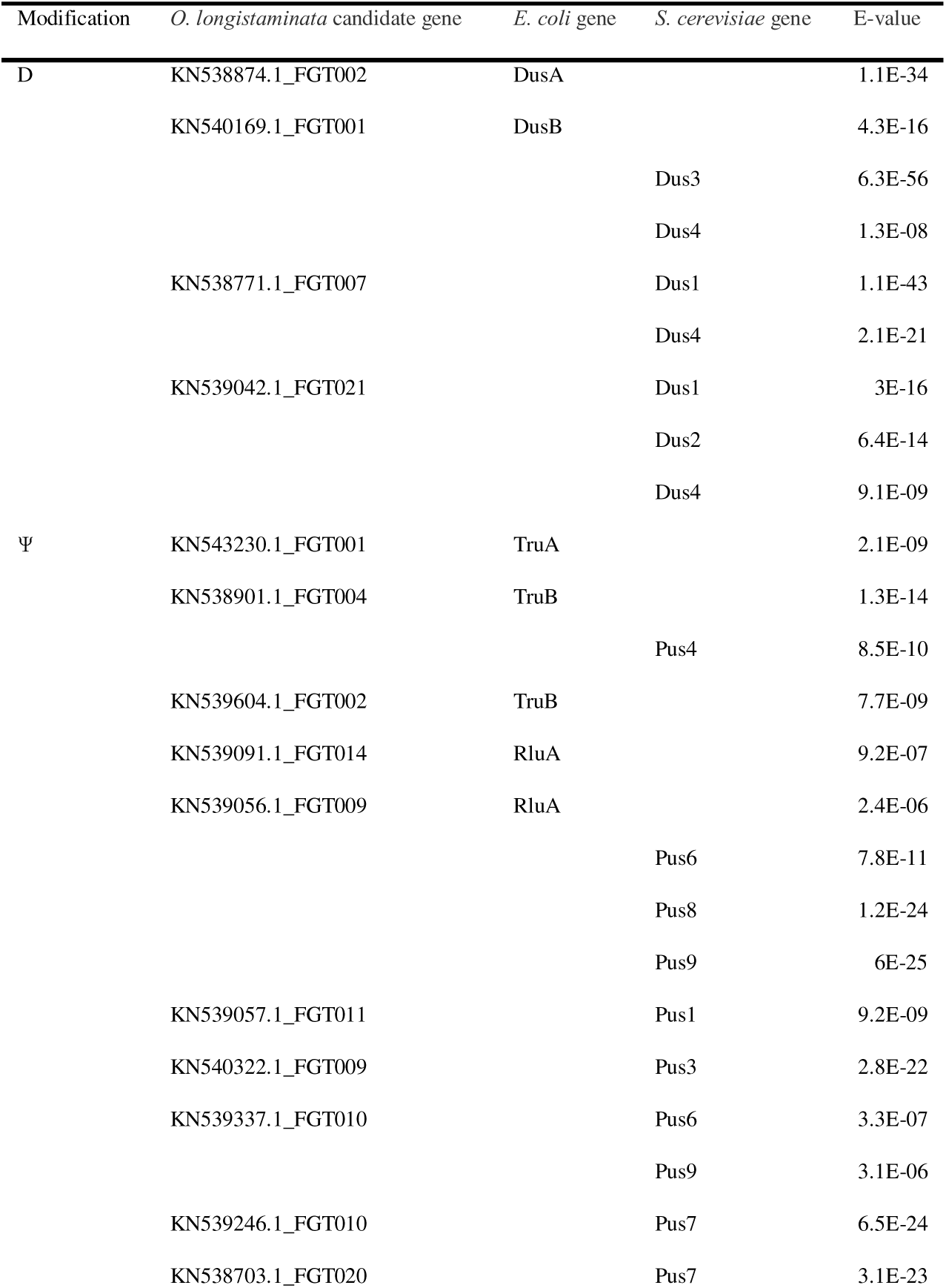
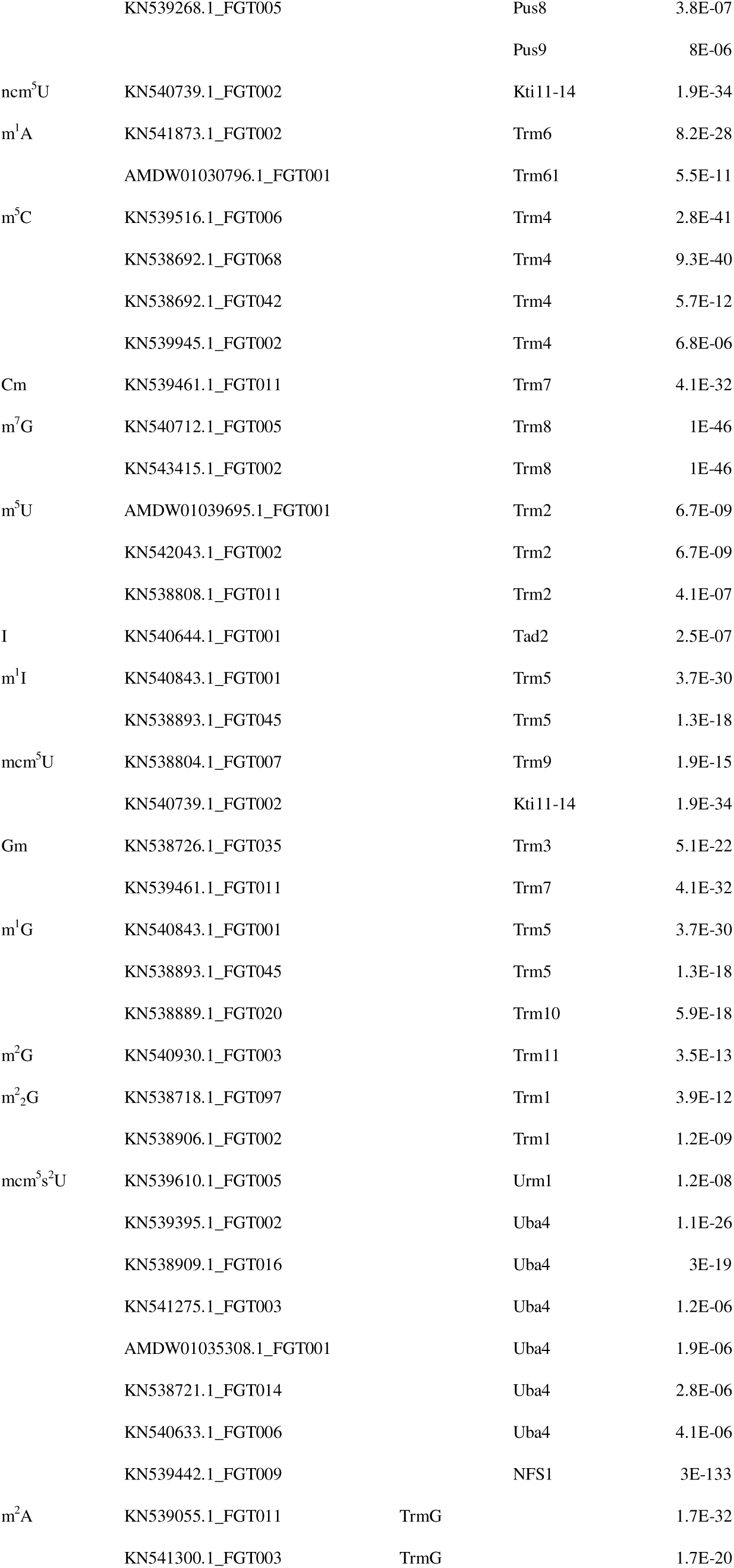
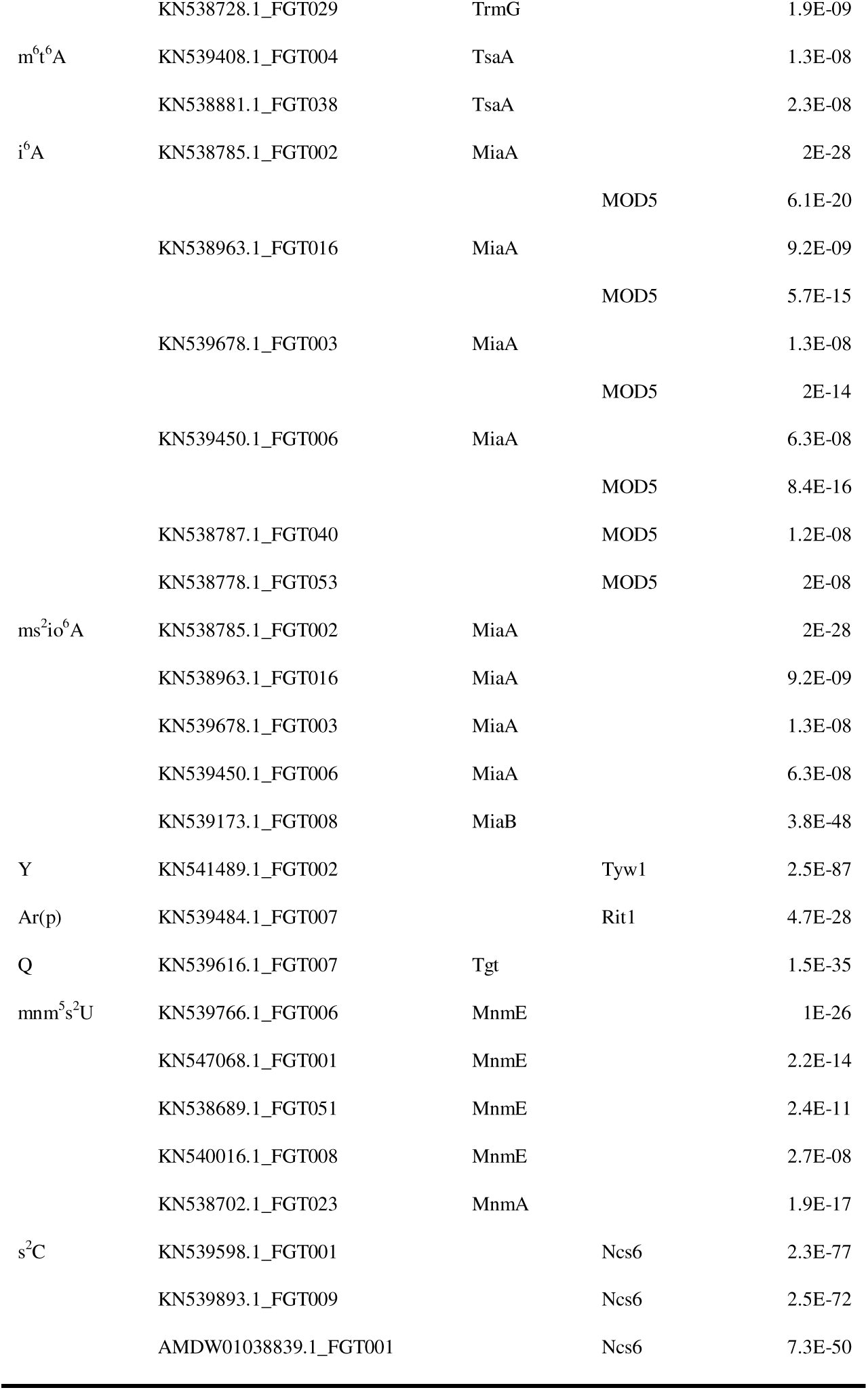
*Oryza longistaminata* candidate genes for tRNA modification.

The expression levels of all tRNA modification candidate genes in *O. longistaminata* are plotted in Figure 5A. Supporting the potential importance of tRNA modification in rhizome development, the majority of the candidate genes were found to be of a higher expression level in rhizomes than in other tissues. Interestingly, except for *KN538787.1_FG040*, *AMDW01030796.1_FG001*, and *KN538963.1_FG016*, none of those candidate genes were detected to be relatively highly expressed in the rhizome elongation zone, suggesting that tRNA modification occurs less intensively in cells undergoing active division and elongation. It is also worth noting that candidate genes responsible for the same tRNA modification can exhibit very different expression profiles. For example, for m^1^I and m^1^G, two modifications that were identified to be important during rhizome development, they shared two corresponding candidate genes—*KN540843.1_FGT001* and *KN538893.1_FGT045*; the former one is mainly expressed in aboveground tissues (the leaf and stem), whereas the latter one’s expression tends to be rhizome-specific. Thus, the expression profiles of the candidate genes might enable us to pinpoint critical genes for the identified key rhizome-related tRNA modifications, all of which have a least one candidate gene showing high expression in the rhizome.

**Figure 5.**
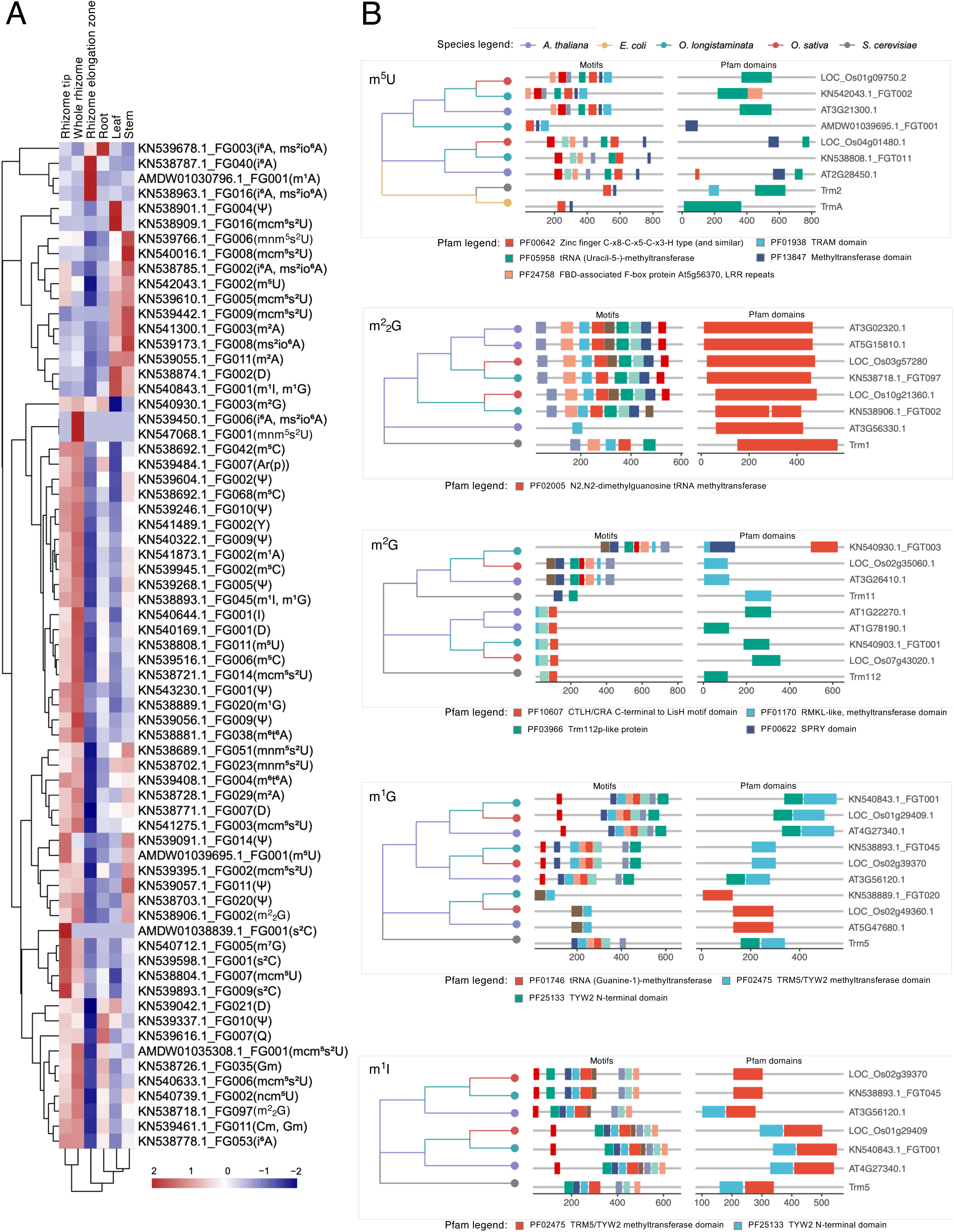
Genome-wide analysis of tRNA modification-related genes in *O. longistaminata*. **A** Expression levels of the tRNA modification candidate genes identified in *O. longistaminata*. The heatmap shows Z-score normalised expression data in five *O. longistaminata* tissues (rhizome tip, whole rhizome, rhizome elongation zone, root, and stem). Clustering was performed using the default Euclidean distance-based clustering algorithm of the ‘pheatmap’ package in R. Corresponding tRNA modifications are shown in the bracket. **B** Phylogenetic tree, motif, and Pfam analysis of genes associated with tRNA modifications. Only differentially behaving modifications with relevance to the underlying hormonal signalling during rhizome development are shown. For all results, see Supplementary Table 1.

To obtain further evolutionary and functional insights for the candidate genes, we constructed a phylogenetic tree for each modification and performed motif and Pfam analyses (Supplementary Table 1). Apart from *O. longistaminata* genes, we also included homologues from *O. sativa*, *A. thaliana*, *E. coli*, and *S. cerevisiae* in these analyses. Visualised in Figure 5B are results for the differentially behaving tRNA modifications with relevance to the underlying hormonal signalling. Overall, candidate genes from *O. longdistaminata* were closely clustered with *O. sativa* genes, harbouring generally conserved motifs and Pfam domains, mainly methyltransferase domains. The only exception was *AMDW01039695.1_FGT001*, an *O. longistaminata* candidate gene for m^5^U. It was found to be phylogenetically closer to one homologue from *A. thaliana*. Pfam search also revealed that some *O. longistaminata* candidate genes, despite showing significant sequence similarity with their homologues, may contain additional domains. For example, an F-box domain and an SPRT domain were located in the m^5^U candidate gene *KN542043.1_FGT002* and m^2^G candidate gene *KN540930.1_FGT003*, respectively, but not in their closest *O. sativa* or *A. thaliana* homologous genes (Figure 5B). Of note, both these domains have reported involvement in signalling pathways (Lechner et al. 2006; Woo et al. 2006), hinting that these *O. longistaminata* candidate genes are more likely associated with rhizome development than other ones in the same phylogenetic tree.

## DISCUSSION

The considerable developmental plasticity of the rhizome system in response to environmental inputs enables rhizomatous plants to achieve highly efficient clonal foraging (Macdonald and Lieffers 1993). To support such a vegetative reproduction strategy, rhizomatous plants supposedly utilise a sophisticated regulation system for swiftly fine-tuning their rhizome development. In non-plant systems, there is a large body of evidence signifying the importance and efficiency of tRNA modifications in developmental regulation (Frye et al. 2018). Two modes of action have been revealed. The first is via tRNA-derived small noncoding RNA fragments (tRFs), whose biogenesis is shaped by the tRNA modification landscape (Chen et al. 2021; Liu et al. 2021). Certain modifications protect tRNAs from cleavage, leading to low intracellular abundances of tRFs and high global translation (Blanco et al. 2014; Blanco et al. 2016), and meanwhile, tRFs could also impair global and gene-specific protein synthesis by displacing RNA-binding proteins from mRNA during the translation process (Sharma et al. 2016; Guzzi et al. 2018). Second, modifications at the wobble position can enhance the recognition versatility of a tRNA molecule to the genetic code in mRNAs, thereby optimising codon usage and translation (Schaffrath and Leidel 2017). While these in-depth mechanisms have not yet been examined in plants, the results of this study suggest that rhizome development is likely also underpinned by the dynamic deposition and removal of modifications in tRNAs. Intriguingly, several rhizome-relevant modifications identified here (e.g., ψ, m^5^C, and m^2^G) have been used to exemplify the above-mentioned mechanisms (Van Haute et al. 2016; Zhang et al. 2018; Guzzi et al. 2018); rhizome development in *Oryza longistaminata* may therefore represent a useful platform for exploring tRNA modification-mediated gene expression regulation in plants.

Nutrition availability and depletion have been uncovered as a critical factor in regulating rhizome development (Fan et al. 2022). When a rhizome elongates, a carbohydrate gradient is gradually formed from the base to the tip of the rhizome, and it is hypothesised that this gradient serves as a developmental cue (Bessho-Uehara et al. 2018). In the present study, we found that the involvement of tRNA modification in rhizome development seemed to be more marked during the middle developmental stage, indicating that the nutritional status may interconnect with the tRNA modification pattern. Indeed, using non-plant models, links have been established between nutrient inputs and tRNA modifications, with energy metabolism as a central hub (Rashad and Marahleh, 2025). While the precise mechanisms remain elusive, it is clear that the nutritional status can influence enzymatic tRNA modifications via regulating the supply of precursors and enzyme substrates (Accornero et al. 2020). In addition to carbohydrates, nitrogen sources in the soil also, to a certain extent, govern the rhizome structure (Shibasaki et al. 2021; Kawai et al. 2022). In this study, we compared two nitrogen conditions, a nitrogen-rich one and a nitrogen-limiting one. Of note, the low-N condition represented a metabolic stress to plants, for a dampened nitrogen metabolism leads to insufficient amino acids. In yeast, cells can use modified tRNAs to perceive amino acid availability, and interestingly, tRNA modification plays a role in balancing carbon and nitrogen metabolic flux for optimal growth (Gupta et al. 2019). Whether such a tRNA-based regulatory system exists in rhizome development is worthy of further research.

The connection between hormonal signalling and tRNA modification is less understood, in particular for phytohormones. In this study, exogenous hormone treatments led to overlapping but broader alterations in the tRNA modification pattern in comparison with the corresponding external environmental factors, despite the fact that they exerted similar effects on the rhizome phenotype. This is not surprising as exogenous hormone treatments could hardly precisely mimic the endogenous situation and may cause a wider spectrum of cellular changes. Nevertheless, the shared key modification types suggest that tRNA modification is presumably at play downstream of hormonal signalling. With little research conducted to study their interplay in plants, perhaps looking at some conserved signalling components across kingdoms may offer some clues. Extensive studies in yeast, animals, and plants have documented target of rapamycin (TOR) protein kinase as a master signalling integrator orchestrating metabolism, growth, and developmental transitions in response to environmental cues (Wu et al. 2019; Liu and Sabatini 2020). Importantly, in non-plant species TOR has been discovered to lie at the nexus of metabolic shifts and tRNA modification dynamics (Rashad and Marahleh 2025), and in plants it coordinates nutrient, energy, light, and hormone signalling (including cytokinin and GA signals) (Marash et al. 2024). Taken together, there is also likely a TOR-tRNA axis in plants, and to test out this hypothesis, functional experiments using tor and hormone biosynthesis mutants are warranted.

## CONCLUSIONS

A deeper understanding of rhizome development is important for further breeding efforts of perennial grain crops, which are promising elements of future sustainable agriculture (Li et al. 2022). This study focused on rhizome-related tRNA modifications. Through comparative profiling analysis, we identified a couple of tRNA modifications that potentially play roles in rhizome’s elongation and developmental transition and foregrounded the interplay between hormonal signalling and tRNA modification in rhizome development. These results, coupled with our genome-wide analysis of tRNA modification-related genes, may lay a foundation for future rhizome biology research and support perennial crop development. More importantly, as an initial attempt to systematically explore tRNA modification in plant organ development, our results provide evidence that in plants tRNA-based translational regulation may echo its developmental importance revealed in various other organisms, and this may open up exciting new avenues to the plant research community.

## MATERIALS AND METHODS

### Plant material and growth conditions

To ensure that rhizome materials were of the same generation, for each experiment a single *O. longistaminata* plant was first grown in a greenhouse under long-day conditions (16-h light/ 8-h dark) at an average temperature of 28 °C, and its rhizomes were excised and cut into sections that each contained a node. These rhizome sections were transferred to standard Yoshida solution in a growth chamber under a 16-h day photoperiod at 28 °C and 60% relative humidity for shooting, and the newly formed ramet was then further hydroponically cultured using an opaque black bucket in the greenhouse under the same conditions for rhizome sample collection.

### Treatments and phenotypic observations

For experiments investigating the elongation stage of rhizomes, hydroponically cultured *O. longistaminata* plants that formed rhizomes with various lengths were transferred to a modified Yoshida solution for treatment, and plants cultured in standard Yoshida solution were used as a control. High nitrogen (high-N), low nitrogen (low-N), and tZ treatments were conducted as previously described (Shibasaki et al., 2021). Elongating rhizomes were categorised into three groups (early stage, middle stage, and late stage) according to the number of internodes. To measure rhizome bud activity, node-containing rhizome sections were implanted in 0.6% agar medium using sterile culture jars, and shoot length was measured after two weeks of culturing.

For experiments investigating the developmental transition stage of rhizomes, elongating rhizomes that curved with an angle less than 15 degrees and harboured 6–8 nodes were excised from the hydroponically cultured *O. longistaminata* plants. For treatments, these rhizomes were *ex vivo* cultured in the dark using a 15 ml Falcon tube containing 1/2 Murashige and Skoog (MS) liquid medium with or without supplements. GA and sucrose treatments were conducted as previously described (Bessho-Uehara et al. 2018), and rhizomes cultured in 1/2 MS medium were used as a control. For light treatment, rhizomes were cultured in Falcon tubes without aluminium foil wrapped, and the photoperiod was set to 16-h light/8-h dark. For all the groups, curvature growth phenotypes were photographed every two days under green light during the eight-day period of culturing, and the angle was measured by using the Fiji platform (Schindelin et al. 2012).

### tRNA isolation

To obtain tRNAs from rhizome samples, a three-step isolation protocol was employed (Chen et al., 2010). Briefly, total RNA was first extracted using Trizol Reagent (Invitrogen, Carlsbad, USA) following the manufacturer’s instructions, and its quantity, integrity and purity were evaluated using agarose electrophoresis and a NanoDrop ND-1000 spectrophotometer (Thermo Fisher Scientific, Waltham, USA). Second, sRNAs (i.e., tRNAs, miRNAs, and snRNAs) were separated from large-size RNA molecules (i.e., rRNAs and mRNAs) by mixing LiCl solution (4 M) with the same volume of total RNA solution. After centrifugation at 8000 r/min at 4 °C for 30 min, the pellet containing rRNAs and mRNAs was discarded, and sRNAs in the supernatant were precipitated with three volumes of ethanol, followed by a washing step with 70% ethanol. sRNAs were dissolved in a solution containing 0.1 M Tris and 0.1 M NaCl (pH = 7.4). Finally, tRNAs were further isolated from the sRNA mixture by using DE52 cellulose anion exchange resin following the protocol described by Chen and Xu (2025) with slight modifications. The sRNA solution was loaded into a pre-prepared DE52 cellulose column and passed through it by gravity at room temperature. Subsequently, the column was washed three times with 5 ml DE52 binding buffer (0.1 M Tris, 0.1 M NaCl, pH = 7.4), and the tRNA molecules were eluted into a 15 ml centrifuge tube using 7 ml tRNA elution buffer (0.1 M Tris, 1 M NaCl, pH=7.4). To precipitate tRNAs, 0.7 volume of isopropanol was added to the tube, and the mixture was incubated at –20 °C overnight. Following centrifugation (8000 r/min, 4 °C, 30 min), the pellet was washed with 70% ethanol, and tRNAs were dissolved in deionised water.

### LC-MS/MS-based nucleoside analysis

To analyse modified tRNA nucleosides, tRNA molecules were first degraded by mixing 20 μg tRNA with 4 μl P1 nuclease (New England Biolabs, Ipswich, USA) and 0.6 μl bacterial alkaline phosphatase (Sigma-Aldrich, St. Louis, USA) in HEPES buffer (final concentration: 20 mmol/l). The digestion mix was incubated at 37 °C for 3 h, and after centrifugation at 12,000 r/min at room temperature for 15 min, degraded tRNA samples were diluted using deionised water to a concentration of 15 ng/μl. The LC-MS/MS analysis was performed as previously described (Wang et al., 2017). In brief, degraded tRNA samples were loaded into a custom setup consisting of an API 4000 Q-Trap mass spectrometer (Applied Biosystems, Waltham, USA), an LC-20A HPLC system equipped with an Inertsil ODS-3 column (2.1 × 150 mm, 5 μm particle size; Shimadzu, Kyoto, Japan), and a diode array UV detector. Electrospray ionisation mass spectrometry was conducted in positive ion mode with the following settings: curtain gas: 15 psi, auxiliary gas: 65 psi, nebuliser gas: 60 psi, turbo gas temperature: 550 °C, entrance potential: 10, and ion spray voltage: 5500 V. The mobile phase was composed by 2 mM ammonium acetate (solvent A) and methanol (solvent B), and the parameters of the elution were set as follows: injection volume: 4 μl; column temperature: 30 °C; and gradient programme: 0% of solvent B and 100% of solvent A during 0 to 10 min, 50% of solvent B to 50% of solvent A during 10 to 13 min, 0% of solvent B and 100% of solvent A during 13 to 23 min, 5% of solvent B and 95% of solvent A during 23 to 23.1 min, and 0% of solvent B and 100% of solvent A during 23.1 to 30 min. Using multiple reaction monitoring mode, a total of 36 tRNA modification types were quantified by setting the following parameters: declustering potential, collision energy, collision cell exit potential, *m*/*z* of the transmitted parent ion, *m*/*z* of the monitored product ion, and retention time. Specific parameter settings of each nucleoside were the same as previously described (Yan et al. 2013). All nucleoside standards were produced by Santa Cruz Biotechnology (Dallas, USA).

### Genome-wide analysis of tRNA modification-related genes

Potential tRNA modification-related genes in *A. thaliana*, *O. sativa*, and *O. longistaminata* were probed by using protein sequences of functionally characterised tRNA-modifying genes in yeast (Chen et al., 2010). For *A. thaliana*, its genes were obtained by using the built-in ‘blastp’ tool of The Arabidopsis Information Resource (http://www.arabidopsis.org) with default settings. For *O. sativa* and *O. longistaminata*, their genome sequences were first downloaded from the Ensembl Plants database (Zhang et al. 2015). Subsequently, TBtools-II was employed to convert coding DNA sequences into protein sequences, and the built-in ‘blast’ and ‘Sequence Toolkit’ functions were used with default settings for obtaining homologous tRNA modification-related genes (Chen et al. 2023). For expression analysis of tRNA-modifying genes in *O. longistaminata*, publicly available RNA-seq data were downloaded from GenBank’s Short Read Archive (accession numbers: PRJNA196977, SRR828682, SRR830212, SRR830652, SRR833549, SRR831108, SRR834501, SRR834502 and SRR831166) (He et al. 2014). The fastp software was employed for raw read preprocessing and quality control (Chen 2023). Clean reads were obtained by removing low-quality reads and those with more than 10% of unrecognisable bases. Alignment of reads to the reference *O. longistaminata* genome was performed using HISAT2 (2.1.0) (Kim et al. 2019), generating positional and sequence information for the samples. The counts of the genes were obtained through the featureCounts program (Liao et al. 2013), and the transcripts per kilobase of exon model per million mapped reads (TPM) were calculated in the R environment. For phylogenetic analysis, multiple sequence alignment, tree construction, and tree visualisation were performed by using MUSCLE5 (Edgar 2022), MEGA12 (Kumar et al. 2024), and the ‘ggtree’ package in R, respectively. Motif identification was conducted using the MEME suite (5.5.8) (Bailey et al. 2015). Pfam domains were scanned by the pfam-scan script (1.0) using the pfam-A database (http://pfam.xfam.org), and the results were visualised using the ‘gggenes’ package in R.

### Statistical analysis

All statistical analyses were performed in the R environment. For multiple comparisons between treatments, one-way or two-way ANOVA was performed with Tukey’s honest significant difference test. For binary comparisons, significant differences were determined by two-tailed Student’s *t*-test. A *P*-value < 0.05 was considered significantly different.

## SUPPLEMENTARY DATA

**Supplementary Table 1.** Phylogenetic tree, motif, and Pfam analysis of genes associated with tRNA modifications.

## ACKNOWLEDGEMENTS

We thank Dr. Friedrich Kragler (Max Planck Institute of Molecular Plant Physiology) for helpful discussions.

## AUTHOR CONTRIBUTIONS

Z.L., P.C., F.H., and X.L. designed and supervised the study. W.L., G.Y., C.Z., and R.Y. conducted the experiments. W.L., G.Y., H.X., R.M., and Y.L. performed the analyses. Z.L., G.Y., and R.M. wrote the manuscript with additional inputs from all the authors. All authors read and approved the paper.

## DATA AVAILABILITY

Data will be made available on request.

